# Prenatal cadmium exposure alters proliferation in mouse CD4^+^ T cells via LncRNA Snhg7

**DOI:** 10.1101/2021.06.04.446938

**Authors:** Jamie L. McCall, Melinda E. Varney, Sebastian A. Dziadowicz, Casey Hall, Kathryn Blethen, Gangqing Hu, John B. Barnett, Ivan Martinez

**Author notes:** Department of Pharmaceutical Science, Marshall University, Huntington, WV 25701, USA. **Correspondence:** Ivan Martinez.

## Abstract

**Objective:** Prenatal cadmium (Cd) exposure leads to immunotoxic phenotypes in the offspring affecting coding and non-coding genes. Recent studies have shown that long non-coding RNAs (lncRNAs) are integral to T cell regulation. Here, we investigated the role of long non-coding RNA small nucleolar RNA hostgene 7 (lncSnhg7) in T cell proliferation.

**Methods:** RNA sequencing was used to analyze the expression of lncRNAs in splenic CD4^+^ T cells with and without CD3/CD28 stimulation. Next, T cells isolated from offspring exposed to control or Cd water throughout mating and gestation were analyzed with and without stimulation with anti-CD3/CD28 beads. Quantitative qPCR and western blotting were used to detect RNA and protein levels of specific genes. Overexpression of a miR-34a mimic was achieved using nucleofection. Apoptosis was measured using flow cytometry and luminescence assays. Flow cytometry was also used to measure T cell proliferation in culture.

**Results:** We identified 23 lncRNAs that were differentially expressed in stimulated versus unstimulated T cells, including lncSnhg7. LncSnhg7 and a downstream protein, GALNT7, are upregulated in T cells from offspring exposed to Cd during gestation. Overexpression of miR-34a, a regulator of lncSnhg7 and GALNT7, suppresses GALNT7 protein levels in primary T cells, but not in a mouse T lymphocyte cell line. The T cells isolated from Cd-exposed offspring exhibit increased proliferation after activation *in vitro*, but Treg suppression and CD4^+^ T cell apoptosis are not affected by prenatal Cd exposure.

**Conclusion:** Prenatal Cd exposure alters the expression of lncRNAs during T cell activation. The induction of lncSnhg7 is enhanced in splenic T cells from Cd offspring resulting in the upregulation of GALNT7 protein and increased proliferation following activation. miR-34a overexpression decreased GALNT7 expression suggesting that the lncSnhg7/miR-34a/GALNT7 is an important pathway in primary CD4^+^ T cells. These data highlight the need to understand the consequences of environmental exposures on lncRNA functions in non-cancerous cells as well as the effects *in utero*.

## 1 INTRODUCTION

Cadmium (Cd) heavy metal exposure is of concern due to its long half-life and its association with numerous health issues including pregnancy and reproductive disorders, developmental toxicity, and cancer (1). Cd exposures occur through cigarette smoke and eating food from crops grown in contaminated soil. Occupational Cd exposures occur during mining, work with Cd-containing ores, and during the manufacturing of products containing Cd such as paints and batteries. The areas surrounding former zinc smelters, such as the site in Spelter, West Virginia, have soil and household dust that is contaminated with extremely high levels (up to 2280 mg/kg) of Cd (SI Group, LP. Dust Sampling Harrison County, West Virginia June and August 2005). The estimated daily intakes of Cd in nonsmoking adult males and females living in the United States are 0.35 and 0.30 μg Cd/kg/day (2), respectively, but reaches higher levels in contaminated areas (3). While there are many routes of human exposure, Cd is a health risk because it does not undergo metabolic degradation to a less toxic product and is poorly excreted (4).

Increasing evidence shows that the developing immune system is particularly susceptible to environmental insult and that exposure during these periods contribute to long-term immune dysfunction in the offspring later in life (5–7). We previously identified numerous immunotoxic effects in offspring of mice exposed to Cd (8–10). At birth (fewer than 12 hours postpartum) (8), prenatal Cd exposure increased the number of CD4^+^ T cells and a subpopulation of double negative cells, DN4 (CD4^−^CD8^−^CD44^−^CD25^−^), indicating that T cell maturation is altered by prenatal Cd exposure. At 2 and 7 weeks of age (9), prenatal Cd exposure did not affect thymocyte populations, implying that the aberrant T cell maturation is not sustained throughout life. However, other sex-specific immune alterations were observed at later ages (10).

Secreted cytokines play an important role in immune cell communication and signaling. Interleukin 2 (IL-2) is secreted from CD4^+^ T cells following T cell receptor stimulation (11) and is also required for the inhibitory activity of regulatory T cells (Tregs) (12). Interferon gamma (IFN-γ) is a signature proinflammatory cytokine in inflammation and autoimmune diseases; however, it also contributes to immune system homeostasis [reviewed in (13)]. Both IL-2 and IFN-γ levels were decreased in male and female prenatal Cd offspring at 7 weeks, but female offspring had reduced IL-2 levels at an earlier age of 2 weeks. By 7 weeks, female offspring exposed to prenatal Cd display a more inflammatory phenotype with reduced Tregs in the spleen. At this age, offspring of both sexes show enhanced T-dependent antibody production in response to immunization with heat-killed *S. pneumoniae* indicating that male Cd offspring were showing increased proinflammatory responses. Increased T-dependent, PspA-specific serum IgG titers remained elevated in both female and male Cd offspring compared to control animals at 20 weeks (10). At 20 weeks, all Cd offspring also had increased proportions of splenic CD4^+^ T cells and CD45R/B220^+^ B cells as well as decreased proportions of CD4^+^FoxP3^+^CD25^+^ (natural regulatory T cells, nTregs) compared to the controls. Taken together, these data demonstrate that prenatal Cd exposure leads to dysregulation of T cell production and function, albeit on different timelines dependent on the sex of the offspring.

Emerging data highlight the integral roles that long noncoding RNAs (lncRNAs) play in T cell function [reviewed in (14)]. LncRNA fragments are greater than 200 nucleotides and lack protein encoding potential (15). They are often times spliced and polyadenylated (16). LncRNAs regulate gene expression through various mechanisms including micro-RNA sponging, histone and chromatin modifications, and protein translation (17, 18). The lncRNA small nucleolar RNA hostgene 7 (lncSnhg7) was first reported in 2013 by Chaudhry et al. (19) in the lymphoblastoid cell lines TK6 and WTK1, which differ only in p53 function. However, lncSnhg7 is aberrantly expressed and has oncogenic properties in a variety of cancers including breast (20, 21), pancreatic (22), hepatic (23), hypopharyngeal (24), and bladder (25–27) cancers, osteosarcoma (28, 29), and neuroblastoma (30). In colorectal cancer patients, high expression of lncSnhg7 is associated with poor prognosis (31) and is associated with dysregulation of N-acetylgalactosaminyltransferase (GALNT family) proteins (31, 32). Here, lncSnhg7 promotes proliferation and metastasis by sequestering miR-216b resulting in the upregulation of GALNT1 (32). Alternatively, lncSnhg7 can sequester the microRNA miR-34a resulting in increased expression of GALNT7 promoting colon cancer progression via PI3K/Akt/mTOR signaling (31). GALNT7 controls the initiation step of mucin-type O-linked protein glycosylation and the transfer of N-acetylgalactosamine to serine and threonine amino acid residues. Though little is understood regarding lncSnhg7 influence on T cell characteristics, it is known that O-linked glycosylation is critical to T cell maturation, trafficking, and survival (33).

It is crucial to determine the mechanisms by which prenatal Cd exposure affects T cells function and the long-term health of the offspring. We focused on CD4^+^ T cells due to our previous data showing altered cytokine profiles and T-dependent antibody production in offspring exposed to prenatal Cd. Our results revealed that a long noncoding RNA, lncSnhg7, was upregulated in primary CD4^+^ T cells of both male and female offspring after *ex vivo* stimulation with anti-CD3/CD28 in a Cd exposure-dependent way. We examined genes identified from the literature whose expression is regulated by lncSnhg7. We found that GALNT7, but not GALNT1, protein correlated with lncSnhg7 expression in activated CD4^+^ T cells. MiR-34a overexpression in primary cells suppressed GALNT7 protein levels. Finally, we found that prenatal Cd exposure increase T cell proliferation in response to exogenous activation. This is consistent with our RNA-seq data which indicate increased expression of genes related to ribosome biogenesis in stimulated T cells from Cd-exposed offspring. We predict that lncSnhg7 functions to sequester miR-34a, thereby controlling the production of GALNT7 and proliferation in T cells. Our findings provide new insights into the functions of lncSnhg7 in T cell activation and possible mechanisms that leading to immune dysfunction in offspring exposed to Cd *in utero*.

## 2 MATERIALS AND METHODS

### 2.1 Reagents

Cadmium chloride (CdCl_2_, C3141) was purchased from Sigma Aldrich and reconstituted at 101.6 mg/L (5000X) in ddH_2_O. Stock CdCl_2_ was diluted to 10 ppm in ddH_2_O, mixed thoroughly, and transferred to water bottles prior to autoclaving.

### 2.2 Mice

Male and female C.Cg-Foxp3tm2Tch/J (006769) and C57BL6/J (000664) mice were purchased from The Jackson Laboratory and housed at West Virginia University in a specific pathogen-free barrier facility with 12h light/dark cycles and fed normal chow (Envigo, 2018 Tekland Rodent Diet). For experiments using Cd-exposed offspring, mice were mated in pairs for 5-7 days with continuous exposure to 10 ppm CdCl_2_ (or unspiked water for the controls) via their drinking water. Female mice were exposed throughout pregnancy. Spiked water was replaced with normal water within 12 hours of the birth of the pups. Control and Cd-exposed offspring were aged to 8-20 weeks. For experiments using mice directly exposed to Cd, male and female mice (8 weeks old) were exposed to CdCl_2_ or unspiked water for 21 days to approximate the duration of exposure of the offspring during gestation.

### 2.3 Single Cell Suspensions

Spleens were harvested from euthanized mice and single cell suspensions prepared as follows. Spleens were dissected immediately after euthanasia and submerged into 5 mL of ice-cold phosphate-buffered saline without calcium or magnesium (PBS; Corning, 21-031-CV). Spleens were homogenized between frosted sections of sterile microscope slides and passed through a 20G needle 3-4 times to obtain single-cell suspensions. Cells were washed once with PBS and red blood cells were lysed using RBC lysis buffer (Sigma-Aldrich, R7757). After neutralization with complete medium [RPMI-1640 medium (Corning, 15-040-CV) supplemented with 10% heat-inactivated fetal bovine serum (FBS, Sigma-Aldrich, F0926), L-glutamine (2 mM; Gibco, 25030-081), HEPES (5mM; Gibco, 15630-080), penicillin/streptomycin (100 units/L and 100 μg/mL; Cellgro, 30-002-C1), and 2-mercaptoethanol (0.05 mM, Sigma-Aldrich, M3148)], cells were washed once in staining buffer [PBS pH 7.5, 0.5% bovine serum albumin (BSA; Sigma-Aldrich, A7030), and 2 mM ethylenediaminetetraacetic acid (EDTA; Sigma-Aldrich, E5134)] for T cell isolation. Viable cells were enumerated using trypan blue and a hemocytometer.

CD4^+^ T cells for all experiments were isolated from total splenocytes using a negative selection kit (Miltenyi, CD4+ T Cell Isolation Kit, mouse). Further purification for RNA sequencing experiments was conducted using FACS sorting. Cells were lysed immediately or stimulated at a ratio of 1:1 with Dynabeads™ Mouse T-Activator CD3/CD28 for T-Cell Expansion and Activation (Gibco, 1152D) for 16 hours (qPCR and RNAseq), 72 hours (proliferation, apoptosis, and western blots), or 5 days (suppression assay). RNA was isolated as described below.

### 2.4 RNA isolation

Purified CD4^+^ T cells were homogenized using the Qiashredder (Qiagen, 79654) and RNA was extracted using the RNeasy Plus Mini Kit (Qiagen, 74134). RNeasy MinElute Cleanup Kit (74204) was used to concentrate or purify the RNA further. Samples were quantified using the NanoDrop One (Thermo Scientific).

For miR-34a qPCR, total RNA was extracted using TRIzol Reagent (Ambion, 15596026) as per manufacturer’s instructions and then treated with Turbo DNA free DNase (Invitrogen, AM1907) for 25 minutes at 37°C. RNA concentrations were determined with a NanoDrop 2000 Spectrophotometer (Thermo Scientific).

### 2.5 RNA sequencing

RNA was quantified using Qubit RNA HS assay (ThermoFisher). The quality was measured on the Agilent 2100 Bioanalyzer using the Agilent RNA 600 Pico Kit. Libraries were prepared at the WVU Genomics and Bioinformatics Core using KAPA mRNA HyperPrep Kit (Roche) using 10 PCR cycles starting with 300-550 ng of RNA. Libraries were ran on the Bioanalyzer with an Agilent HS DNA assay for quality assurance prior to sequencing. RNA sequencing was conducted by Admera Health (South Plainfield, NJ) on the Hi-seq 2×150 platform. Sequencing reads were aligned to the mouse reference genome (mm10) using subread (34). Quantification of reads on transcripts annotated by RefSeq was done with feature counts (35). Transcripts annotation of lncRNAs are from ENCODE (v5); genes with names starting with “Gm” or ending with “Rik” were excluded as their functions are not annotated. Differentially expressed lncRNAs in CD4^+^ T cells upon CD3/CD28 stimulation were identified by EdgeR3 (36) with thresholds of FC > 2, FDR < 0.01, and log_2_ (count per million) > 1. Gene set enrichment analysis against KEGG pathways from MSigDB was carried out with GSEA (37).

### 2.6 RT-qPCR

Reverse transcription was performed using iScript™ Reverse Transcription Supermix for RT-qPCR (Bio-Rad, 170-8840) with 100 ng of total RNA per 20 μL reaction. RT-qPCR was performed using the primers and conditions listed in Table 1. All targets were amplified using SsoAdvanced™ Universal SYBR Green Supermix (Bio-Rad) with 40 cycles of a 2-step program (95°C × 10 sec, T_m_ × 30 sec for mGALNT7, mGALNT1, and mβactin or 95°C × 30 sec, T_m_ × 30 sec for mSnhg7) on a StepOnePlus Real Time PCR system (Applied Biosystems). Data were normalized to βactin. Analysis was performed according to the q-base protocol, as previously published (38, 39).

**Table 1.**
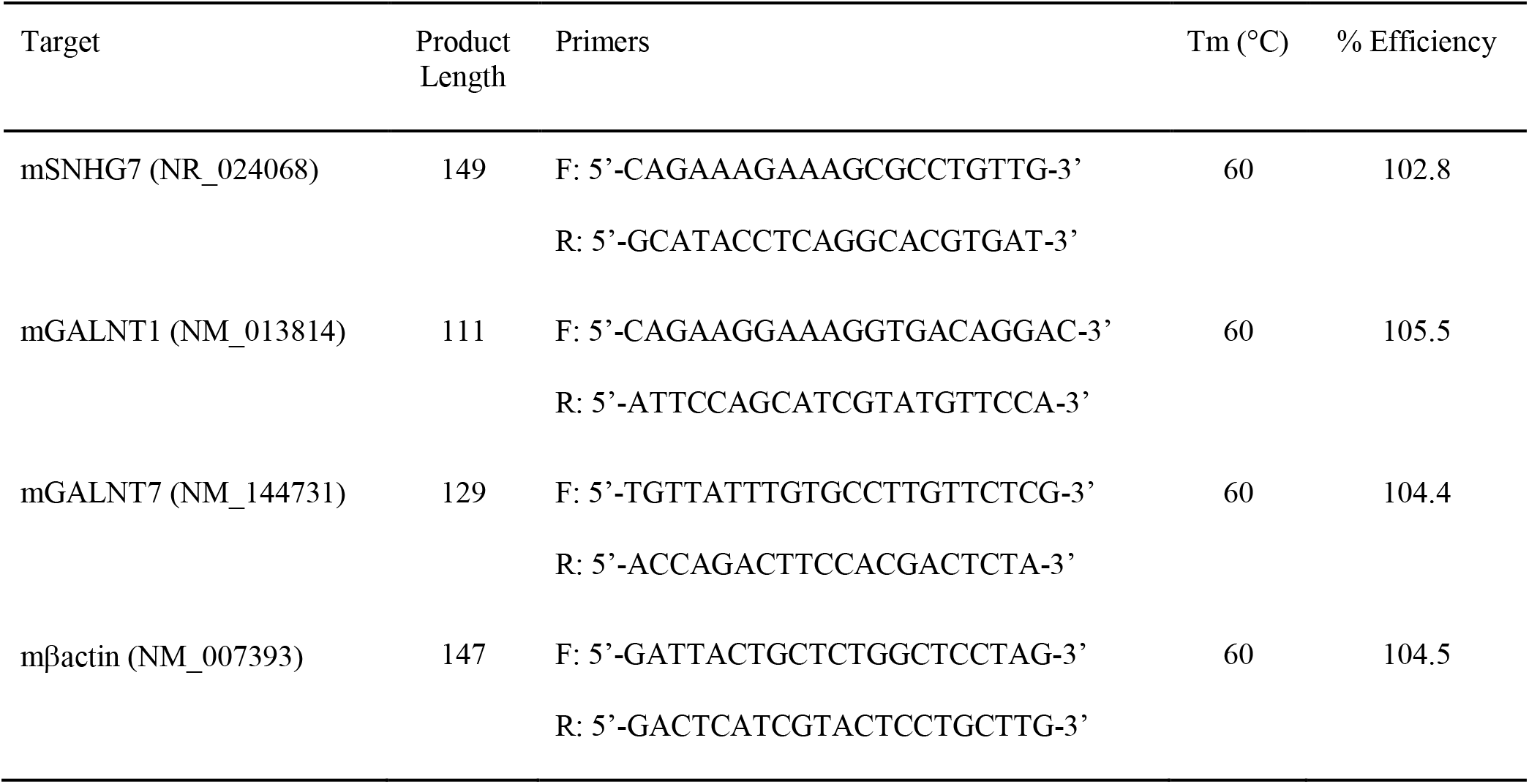
Primers for qPCR

For miR-34a qRT-PCR, 0.5-1 μg of total RNA was reverse transcribed using the TaqMan^®^ miRNA Reverse Transcription Kit (Life Technologies), followed by specific TaqMan^®^ miR-34a Assay (Life Technologies) according to the manufacturer’s protocol. Relative expression was calculated using the ΔΔCt method [relative expression. 2-ΔCt; where ΔCt. Ct (Target RNA) – Ct (endogenous control RNA)], where the endogenous control for miRNA was RNU43.

### 2.7 Nucleofection

Primary CD4^+^ T cells were isolated from mice splenocytes as described above. Cells were nucleofected using the Lonza Amaxa P3 Primary Cell 4D Nucleofector Kit L (V4XP-3024, lot F-13316) according to the manufacturer’s protocol. Briefly, 30, 60, or 90 pmol of Cy3 negative control #1 (AM17120, lot ASO0JHC5), negative control #1 (4464058, lot ASO2F9B3), or miR-34a mimic (MC11030, lot ASO2GN0S) was added to cuvettes containing 1×10^6^ cells and pulsed using program DN-100.

EL4 (ATCC TIB-39) cells were grown in RPMI complete medium. Cells were nucleofected using the Lonza Amaxa SF Cell Line, 4D Nucleofector Kit L (V4XC-2024, lot F-13689) according to the manufacturer’s protocol. Briefly, 90 pmol of negative control #1 (4464058, lot ASO2F9B3) or miR-34a mimic (MC11030, lot ASO2GN0S) was added to cuvettes containing 1×10^6^ cells and pulsed using program CM-120.

For both cell types, cells were incubated for 2 hours prior to stimulation with ratio of 1:1 with Dynabeads™ Mouse T-Activator CD3/CD28 for T-Cell Expansion and Activation (Gibco, 1152D) for 48 hours. RNA and protein were isolated as described above and below, respectively.

### 2.8 Western Blots

Whole cell lysate extracts were prepared M-PER Mammalian Protein Extraction Reagent (Thermo Scientific, 78503) supplemented with 1X Halt Protease and Phosphatase Inhibitor Cocktail (Thermo Scientific, 78447). Protein concentration was determined using the Pierce BCA Protein Assay Kit (Thermo Scientific, 23225). SDS-PAGE was performed using the Bolt system (Invitrogen) including 1X LDS sample buffer and reducing agent, 4-12% Bis-Tris Plus gels, and MES running buffer. Proteins were transferred to PVDF membranes using the iBlot2 transfer system (Invitrogen). Membranes were blocked in Superblock T20 (PBS) Blocking Buffer (Thermo Fisher, 3716) and incubated in primary antibody (listed below) in 5% BSA in TBST overnight at 4°C. Secondary antibodies (listed below) were diluted in 0.1% TBST. Membranes were imaged using the Bio-Rad ChemiDoc Imager and bands were quantified using Image Lab software (Bio-Rad).

Primary antibodies were diluted as follows: GALNT1 (NBP1-81852, lot A115764, Novus) 1:1000; GALNT7 (NBP2-39021, lots R89578 and 000010086, Novus) 1:1000, GAPDH (D16H11; 51745, lot 4, Cell Signaling Technologies) 1:1000, Actin (Ab-5, 612652, lot 6176513, BD Transduction Laboratories) 1:10,000. Secondary antibodies: Goat anti-Mouse Ig (554002, lot 5247553, BD Pharmingen) 1:10,000, Goat anti-Rabbit IgG (15015, lot 04318009, Active Motif) 1:10,000.

### 2.9 Proliferation

Proliferation was assessed using Cell Trace Violet (405/450), (Invitrogen, CellTrace™ Violet Cell Proliferation Kit, for flow cytometry, C34557). Briefly, CD4^+^ T cells were isolated and stimulated as described above. Cells were washed 1X in PBS and anti-CD3/CD28 beads were removed using a magnet. Cells were fixed for 30 minutes at room temperature in 4% paraformaldehyde (PFA) in PBS (Alfa Aesar, J61899). Cells were washed in PBS, resuspended in 300 uL PBS, and transferred to FACS tubes. Cells were assessed by flow cytometry (BD Fortessa using V450 and FITC/GFP filters). Proliferation was analyzed using FCS Express 6. The proliferation index indicates the of average number of cells that any initial cell became (regardless of its proliferation status) while the division index is the average number of cells resulting from a single dividing cell (removes non-dividing cells from the analysis). GFP-positive cells (T regulatory cells) were negligible but gated out of the analysis. T cells from a C57BL/6 mouse were used for the no stain and Cell Trace Violet single-stained controls. An aliquot of whole splenocytes from C.Cg-Foxp3tm2Tch/J mice were used for the FITC/GFP single-stained control.

### 2.10 Apoptosis

Annexin V and 7AAD staining was performed using eBioscience™ Annexin V Apoptosis Detection Kit eFluor™ 450 (Invitrogen, 88-8006-72). Briefly, cells were washed 1X in PBS (1mL) and the anti-CD3/CD28 beads were removed using a magnet. Cells were washed an additional time in 1 mL of binding buffer and resuspended in 100 uL of binding buffer after centrifugation at 500 RCF × 5 minutes. Annexin V-V450 (5 uL) was added to each tube and cells were incubated at room temperature for 15 minutes. Cells were washed in 1X binding buffer and resuspended in 200 uL of binding buffer. 7AAD (5 uL) was added to each sample tube and samples were analyzed by flow cytometry (BD Fortessa using V450, FITC/GFP, and PE-Texas Red filters) within 4 hours. GFP-positive cells (T regulatory cells) were negligible but gated out of the analysis. T cells from a C57BL6 mouse were used for the no stain and single-stained controls. An aliquot of whole splenocytes from C.Cg-Foxp3tm2Tch/J mice were used for the FITC/GFP single-stained control.

Caspase 3/7 activity was measured using Caspase-Glo^®^ 3/7 Assay (Promega, G8090) per the manufacturer’s protocol. Briefly, 100 uL of stimulated cells (5 replicates per sample) were transferred to a white 96-well plate (Costar, 3917). Caspase-Glo reagents (100 uL) were added to each well, samples were mixed, and incubated at room temperature for 1 hour. Luminescence was recorded for 1000 ms using a SpectraMax iD3 Multi-Mode Microplate Reader (Molecular Devices). Control wells included blanks (cell media plus Caspase-Glo Reagent) and negative controls (T cells from a C57BL6 mouse + Caspase-Glo Reagent).

### 2.11 Suppression Assay

Tregs were isolated from 20-week-old cadmium and control offspring using the Miltenyi Mouse Treg isolation kit (130-091-041, lot 5191010723). Tregs were diluted to concentration of 1 × 10^5^ per mL in RPMI complete medium. A C57BL/6 mouse was used to isolate CD4^+^ Tconv cells. Tconv were stained with Cell Tracer Violet (CT Vio, Invitrogen, C34557, lot 1811777) at a dilution of 1:1000, according the manufacturer’s protocol. Tconv cells were resuspended in RPMI complete media at a concentration of 1 × 10^5^ per mL prior to plating. Cells were plated at ratios of 1:20, 1:10, 1:5, 1:2, and 1:1 as shown in Table 2 and stimulated at a ratio of 1:1 with Dynabeads™ Mouse T-Activator CD3/CD28 for T-Cell Expansion and Activation (Gibco, 1152D). After 5 days, cells were harvested, stained with CD4 (BD Pharmingen™ Alexa Fluor^®^ 700 Rat Anti-Mouse CD4, clone RM4-5) and CD25 (BD Pharmingen™ PE Rat Anti-Mouse CD25), and fixed in 4% PFA. Samples were analyzed on the BD Fortessa, acquired with BD FACS DIVA 8.0, and analyzed using FCS Express 6.0 (De Novo Software). To analyze proliferation/division, cells were gated on CD4^+^, CD25^−^, and FoxP3(GFP)^−^ populations. Proliferation was modeled using the following parameters: Starting Generation = geometric mean of T=0 sample (C57BL/6 T cells, stained with CT Vio and fixed prior to stimulation); Background = geometric mean of non-stained control (C57BL6 T cells, no CT Vio, fixed after proliferation); Max Number of peaks = 10 (females) or 9 (males); Software fitted CV and Peak Ratio.

**Table 2.** Suppression assay

|  | Treg:Teff |  |  |  |  |
| --- | --- | --- | --- | --- | --- |
|  | 1:20 | 1:10 | 1:5 | 1:2 | 1:1 |
| Tconv (μL) | 100 | 100 | 100 | 100 | 100 |
| Tregs (μL) | 5 | 10 | 20 | 50 | 100 |
| Media (μL) | 95 | 90 | 80 | 50 | 0 |
| AntiCD3/CD28 (μL) | 12.5 | 12.5 | 12.5 | 12.5 | 12.5 |

### 2.12 Statistical Analysis

*P* values were calculated using Prism Software (GraphPad, La Jolla, CA). A *p* value of less than 0.05 was considered statistically significant. Values presented here are shown as mean +/− standard deviation (SD) unless otherwise noted.

## 3 RESULTS

### 3.1 IncSnhg7 is upregulated in mouse CD4^+^ T cells following activation

We sought to identify pathways in T cells that are altered due to prenatal Cd exposure using a route of administration and dose of Cd that is consistent with our previous publications (8–10). Male and female mice were exposed to CdCl_2_ in their drinking water at a concentration of 10 ppm or administered normal water for 3 days prior to mating. Mice were mated in pairs to ensure that each litter represented an independent exposure and biological replicate. After 5-7 days of cohabitation, male mice were removed from the mating cage and females remained on their respective treatments throughout gestation (Figure 1A). All water was replaced with unspiked water within 12 hours of the birth of the offspring. The offspring were aged to 8-20 weeks of age for assays. Our previous publications show that at these ages, mice exposed to prenatal Cd have aberrant T cell function (9, 10), including increased T-dependent antibody secretion, as well as altered splenic T cell populations (10).

**Figure 1.**
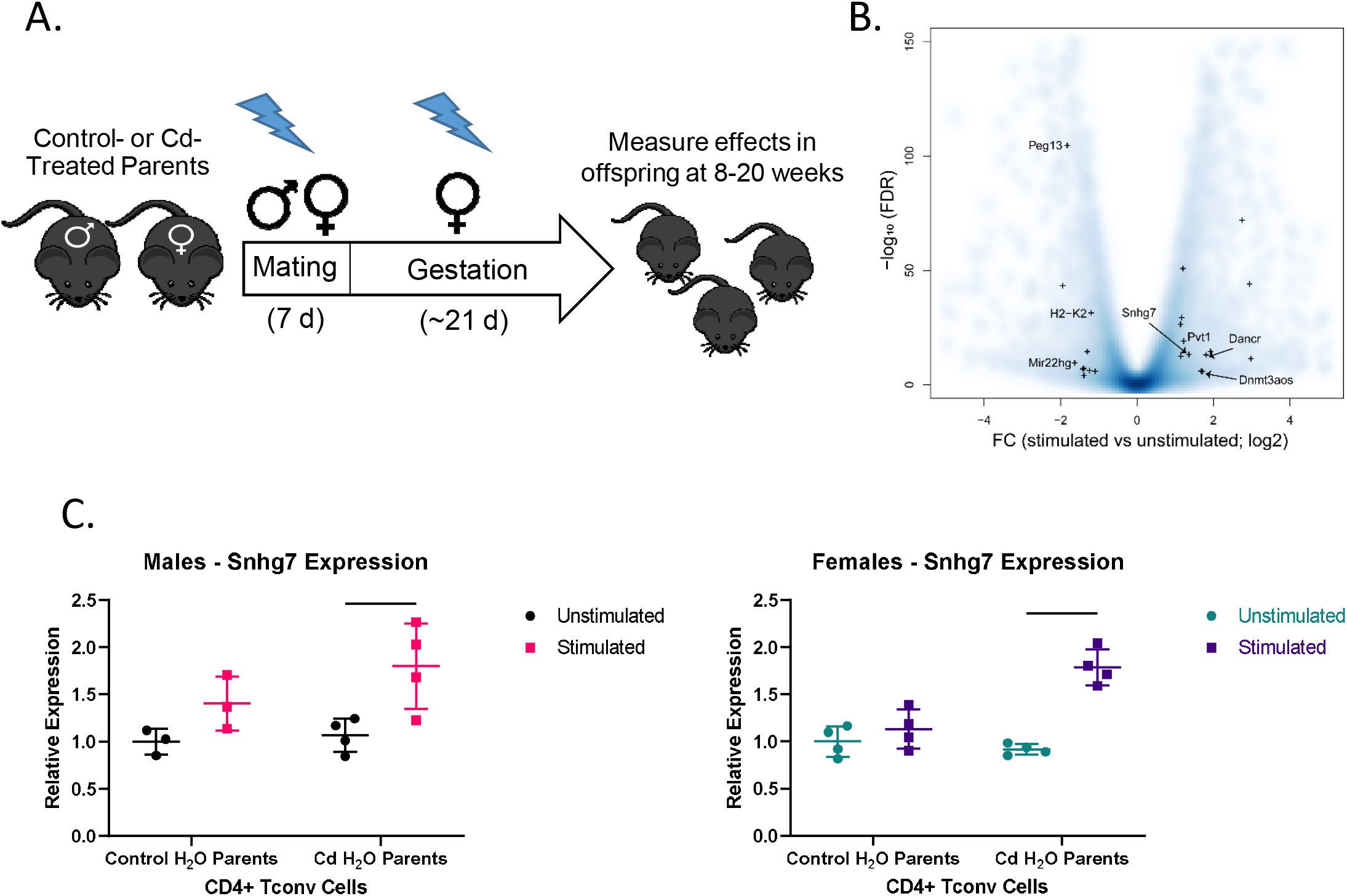
lncSnhg7 expression is increased in CD4^+^ T cells with activation. A) Mice were exposed to Cd during prenatal development. Lightning bolts indicate times where parents are administered CdCl_2_ (10 ppm) via drinking water. B) CD4^+^CD25^−^T conventional (T conv) cells were isolated from the total splenocytes of control offspring. T conv cells were cultured for 0 or 16 hours in the presence of anti-CD3/CD28 magnetic beads, unstimulated and stimulated, respectively. RNA was isolated and expression was analyzed by RNAseq. Volcano plot for the comparison of gene expression between unstimulated and stimulated CD4+ T cells. Blue background represents all genes (coding and non-coding). “+” indicates lncRNAs detected as differentially expressed in our system (FC > 2 and FDR < 0.01). C) CD4^+^CD25^−^ T conventional (T conv) cells were isolated from control and Cd-exposed offspring and stimulated as in B. LncSnhg7 expression was evaluated by qPCR. Statistical significance was assessed using one-tailed, paired t-test between stimulated and unstimulated samples. n=2-4 per group.

The increasing data suggesting that lncRNAs play an important role in T cell function [reviewed in (14)] led us to focus on lncRNAs that are altered in CD4^+^ T cells in our mouse model. CD4^+^ T cells were isolated from total splenocytes of control offspring and stimulated in culture for 16 hours prior to analysis by RNA sequencing. With a modest sequence depth of about 30-million of RNA fragments per library, we identified 23 lncRNA genes differentially expressed in CD3/CD28 stimulated CD4^+^ T cells as compared to unstimulated CD4^+^ T cells (Figure 1B and Table S1). Many of the differentially expressed lncRNAs, including Map2k3os (40), Dnmt3aos (41), AI506816 (42), and Rab26os (43), are relatively uncharacterized; others, such as Snhg4 (44, 45), are only more recently characterized in the literature. While evaluating other lncRNAs in T cell function is of great interest in future studies, we selected lncSnhg7 due its roles in proliferation, apoptosis, and cell differentiation in a variety of cancer cell types (20–30), but a lack of research in primary normal immune cells.

To validate our RNA sequencing results as well as assess the role of prenatal Cd exposure on lncSnhg7 expression, we measured lncSnhg7 in the T cells of control and Cd-exposed mice offspring using qPCR. We observed a general increase in lncSnhg7 expression upon T cell activation; however, the induction of lncSnhg7 expression was only statistically significant in the Cd-exposed offspring (Figure 1C). These results imply that increased lncSnhg7 expression results from T cell activation, but that prenatal Cd exposure further increases lncSnhg7 expression in mice.

### 3.2 GALNT7 protein, but not mRNA, levels are increased in CD4+ T Cells following activation

LncSnhg7 expression affects the protein expression of Polypeptide N-Acetylgalactosaminyltransferase (GalNAc transferase; GALNT) proteins, GALNT1 and GALNT7 (31, 32). GALNT1 and GALNT7 are differentially regulated by lncSnhg7 via its ability to sequester miRNAs, miR-34a and miR-216b, as depicted in Figure 2A. To examine whether lncSnhg7 expression correlates with GALNT levels in mouse CD4^+^ T cells, we measured GALNT1 and GALNT7 mRNA and protein expression with and without activation by anti-CD3/CD28. GALNT1 and GALNT7 mRNA expression was affected by neither prenatal exposure (control or Cd) nor T cell activation (Figure 2B). In contrast, when we measured protein levels, we found that GALNT7, but not GALNT1, was consistently upregulated in CD4^+^ T cells isolated from control and Cd offspring after 72 hours of activation (Figure 2C). In male mice, upregulation of GALNT7 was only significant in the Cd-treated animals, whereas, in the females GALNT7 protein expression was increased regardless of prenatal exposure. These results indicate that there may be a sex-dependent mechanism by which GALNT7 expression is regulated. Additionally, these data indicate that GALNT7 expression is regulated by an Snhg7-independent mechanism as well.

**Figure 2.**
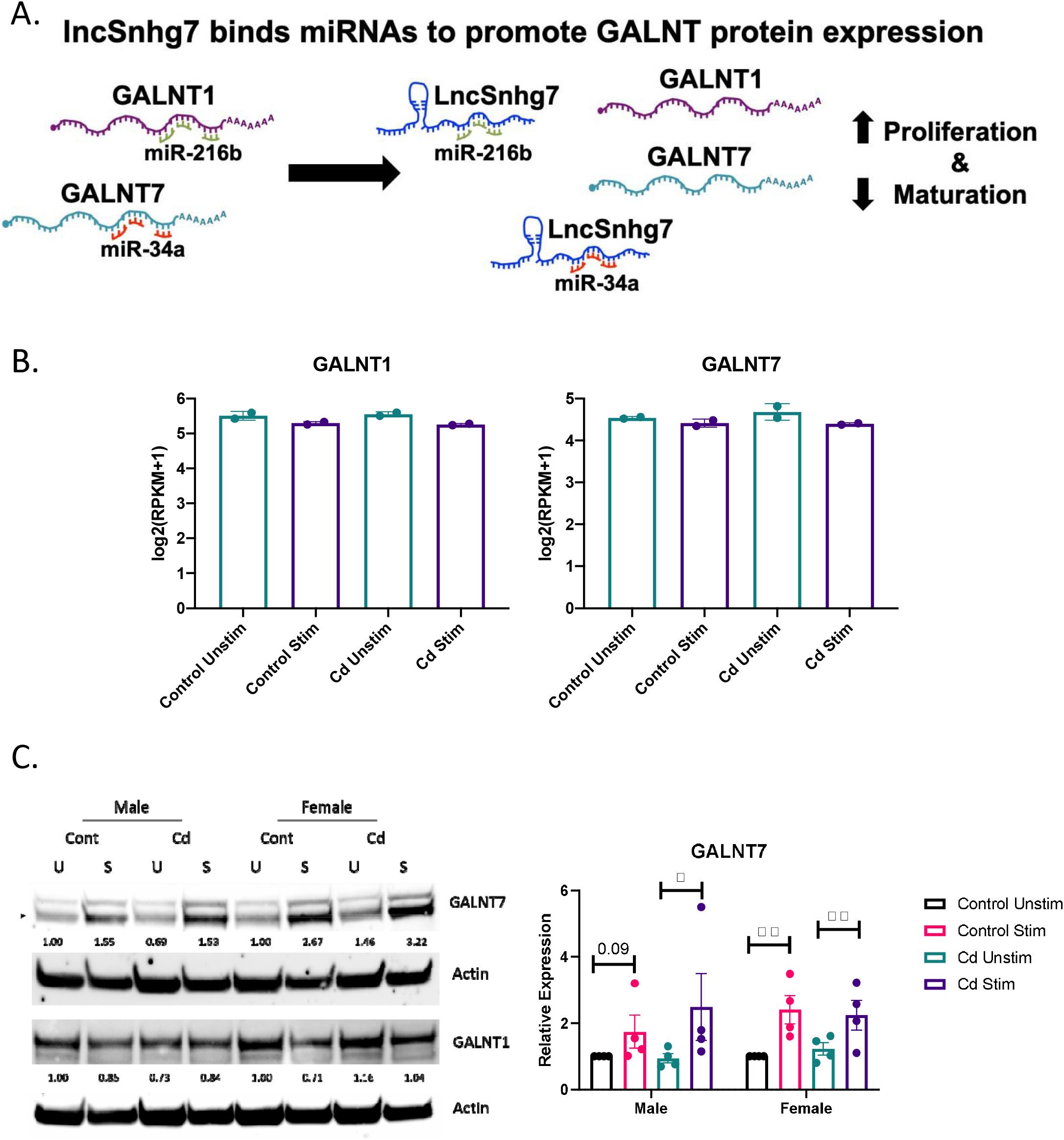
GALNT7 protein expression, but not mRNA expression, is altered in CD4^+^ T cells following activation. A) Model of GALNT1 and GALNT7 regulation by lncSnhg7. B & C) CD4^+^CD25^−^ T conventional cells were isolated from total splenocytes. Cells were cultured in the presence of anti-CD3/CD28 magnetic beads for 0 and 16 hours for RNA or 0 and 72 hours for protein. GALNT1 and GALNT7 expression was analyzed by RNAseq (B) and western blot (C). Statistical significance was assessed using a ratio-matched, one-tailed, paired t-test between stimulated and unstimulated samples.

### 3.3 miR-34a regulates GALNT7 protein expression in CD4^+^ T Cells

The lncSnhg7/miR-34a/GALNT7 relationship has been reported in other cells (31), but has not been established in T cells. To determine whether GALNT7 expression is dependent upon miR-34a levels in T cells, we overexpressed miR-34a using a microRNA mimic. Optimization of miRNA nucleofection was critical to assessing downstream function. We used a Cy3-labeled negative control to measure miRNA uptake using flow cytometry. In primary T cells, 90 pmol of miRNA per 1×10^6^ cells resulted in the brightest median florescent intensity (MFI) per cell (Figure 3A). Additionally, 82.3% of single cells and 31.8% of all events had incorporated the Cy3-labeled microRNA in the 90 pmol sample; whereas only 70.7% single cells (30.4% all events) and 40.8% single cells (12% of all events) were positive for the labeled miRNA in the 60 pmol and 30 pmol groups, respectively. We used 90 pmol per 1×10^6^ cells in the subsequent experiments.

**Figure 3.**
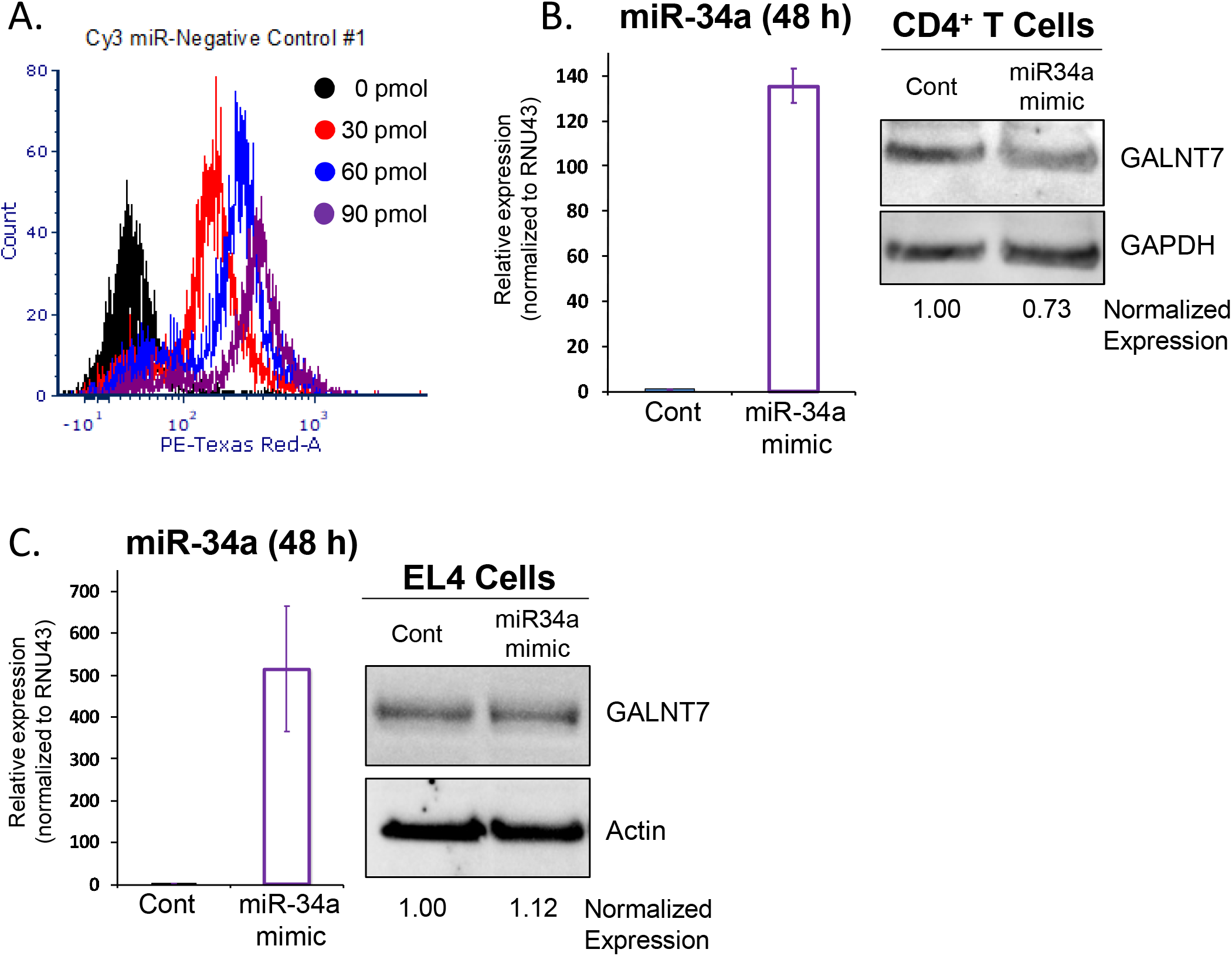
miR-34a regulates GALNT7 expression in primary T Cells. A) Optimization of miR-34a used in nucleofection. miR-34a was overexpressed in mouse CD4^+^ T cells (B) or EL4 cells (C). miR-34a mRNA levels were assessed using qPCR. GALNT7 levels were measured using western blots. Data from representative experiments are shown.

The effects of miR-34a overexpression on GALNT7 protein were assessed in mouse primary CD4^+^ T cells and the mouse T lymphocyte cell line, EL4 (ATCC TIB-39). Of note, both cell types were isolated (or established) from a C57BL/6 mouse strain. In primary CD4^+^ T cells, miR-34a expression was increased 135-fold as compared to the negative control and corresponded with a 27% decrease in GALNT7 expression (Figure 3B) suggesting that miR-34a does regulate GALNT7 expression in T cells. In the mouse EL4 cell line, miR-34a was overexpressed ~500-fold; however, overexpression did not suppress GALNT7 expression (Figure 3C). These results indicate that the miR-34a/GALNT7 relationship may be dysregulated in lymphoma and highlights the importance of experiments using primary cells.

### 3.4 Direct exposure to Cd increases lncSNHG7 and GALNT7, but not GALNT1, expression

All assays described thus far were conducted in cells from offspring that were exposed *in utero* to normal or Cd-spiked water. Previous reports show that while Cd concentrates in the placenta, little to no Cd crosses the placental barrier (46, 47). Therefore, the offspring are not directly exposed to Cd *in utero*. To test whether direct exposure to Cd results in similar effects on lncSnhg7 and GALNT expression, adult male and female mice were exposed to unspiked or Cd water for 21 days. This timepoint was selected as it is the average gestational time of a mouse pregnancy and therefore comparable the prenatal exposure timepoints. CD4^+^ T cells were isolated as described above, stimulated in culture for 0 or 16 hours with anti-CD3/CD28 beads, and mRNA expression was measured by qPCR. LncSnhg7 expression was increased in male and female mice following stimulation regardless of exposure (Figure 4A). In contrast to the data in the offspring where no differences were observed, mice directly treated with Cd exhibited altered expression of GALNT1 and GALNT7 mRNA levels in the T cells following stimulation (Figure 4B and 4C). GALNT7 mRNA levels were increased in the control animals of both sexes, whereas GALNT1 mRNA expression was only increased in the control males (Figure 4B). The changes observed in lncSnhg7, GALNT7, and GALNT1 mRNA levels did not correlate with cadmium exposure and therefore cannot explain the Cd-dependent changes that were observed in the offspring.

**Figure 4.**
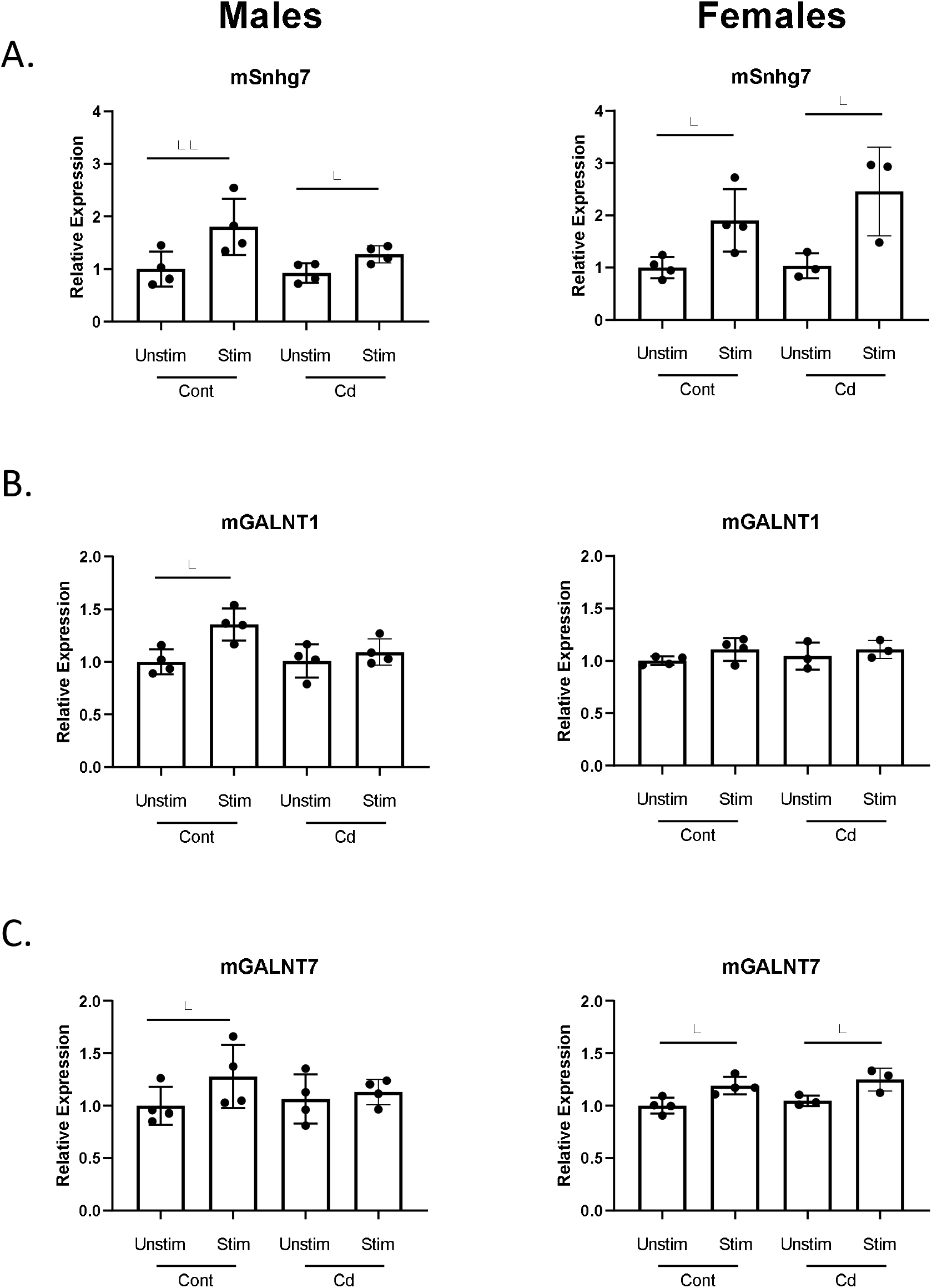
Direct exposure to Cd does not reproduce effects of prenatal Cd exposure. Male and Female mice (8 weeks) were exposed to 21 days of CdCl_2_ via drinking water. CD4^+^CD25^−^T conventional cells were isolated from total splenocytes. Cells were cultured in the presence of anti-CD3/CD28 magnetic beads for 0 and 16 hours prior to RNA isolation. lncSnhg7 (A), GALNT1 (B), and GALNT7 (C) expression was analyzed by qPCR. Statistical significance was assessed using a one-tailed, paired t-test between stimulated and unstimulated samples.

### 3.5 Prenatal cd exposure increases proliferation of CD4^+^ T cells in offspring

LncSnhg7 expression is increased upon stimulation in Cd offspring and we sought to access the effects of prenatal Cd exposure on CD4^+^ T cell function. Previous studies indicate that increased expression of lncSnhg7 and/or GALNT7 regulates cell proliferation in glioma cells (48), cervical cancer HeLa and Caski cells (49), and colon cancer cell lines (31). Therefore, we measured cell proliferation in the CD4^+^ T cells from control and Cd offspring. CD4^+^ T cells were isolated from the spleens of offspring, labeled with Cell Trace Violet, and stimulated in culture with anti-CD3/CD28 beads for 72 hours. Dilution of the Cell Trace Violet was assessed by flow cytometry and proliferation populations were modeled using FCSExpress software. Figure 5A shows examples of individual proliferation plots as well as the quantification of the Division Index (DI) for each mouse. Proliferation was significantly increased in both male and female offspring exposed to prenatal Cd, as indicated by the increased mean DI. T cells from prenatal Cd exposed male offspring had a mean DI of 12.01 ± 2.004 as compared to the controls (3.123 ± 0.032), p<0.01. Female Cd offspring also exhibited increased proliferation (DI = 16.16 ± 4.313) as compared to controls (DI = 9.913 ± 0.6235), p<0.05. Concordantly, at genome-wide transcription level, gene set enrichment analysis against KEGG pathway revealed that ribosome-associated genes were preferentially upregulated in the stimulated T cells isolated from the prenatal Cd-exposed offspring as compared to those isolated from control offspring (Figure S1); ribosome biogenesis is a canonical hallmark of cell proliferation (50).

**Figure 5.**
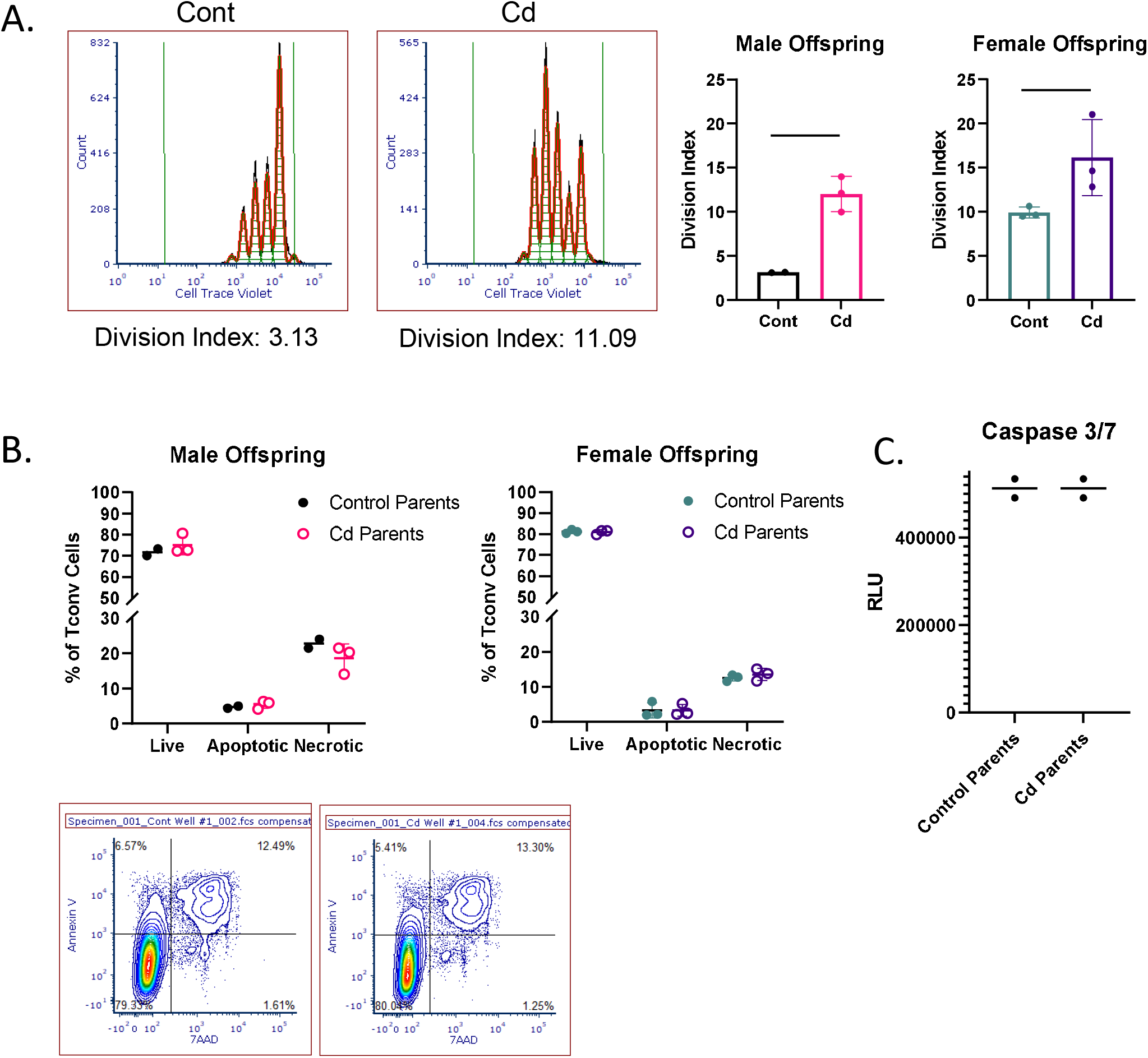
Activated T Cells from Cd offspring have increased proliferation as compared to control offspring, but no differences in apoptosis. A) Splenic CD4+CD25-T l cells were labeled with Cell Trace Violet and stimulated with anti-CD3/CD28 magnetic beads for 72 h. Division index was calculated using FSC Express 6. A one-tailed, unpaired t-test was used to compare groups (n=2-3 per group); * p<0.05, **p<0.01. B & C) Unlabeled T conventional cells were stimulated for 72 hours. Apoptosis was assessed by Annexin V/7AAD staining (B) and caspase 3/7 activity (C).

We have previously reported that prenatal Cd exposure decreases proportions of splenic nTregs at 20 weeks (10). To test whether the increase in proliferation observed here is due to decreased Treg function, we performed a suppression assay using combinations of Treg: Teff (CD4^+^ T Cells) of 1:20 to 1:1. The Tregs of control and Cd offspring suppressed CD4^+^ T cell proliferation equally (Figure S2). To determine if CD4^+^ T cells from Cd offspring exhibited decreased apoptosis, we stimulated cells for 72 hours in culture (without Cell Trace Violet staining) and measured apoptosis using two methods: 1) Annexin V and 7AAD staining was measured by flow cytometry (Figure 5B) and 2) Caspase 3/7 activity was measured using a luminometer (Figure 5C). Both methods suggest that apoptosis was not altered in Cd offspring as compared to control offspring. Taken together, these results indicate that prenatal Cd exposure enhances CD4^+^ T cell proliferation but does not inhibit apoptosis or nTreg inhibitory function.

## 4 DISCUSSION

To our knowledge, we are the first to report that lncSnhg7 is upregulated during CD4^+^ T cell activation in a parent and offspring animal model. We further demonstrate that prenatal Cd exposure exacerbates this phenotype resulting in aberrant T cell proliferation. Several reports in various cancer cells demonstrate that lncSnhg7 regulates cellular proliferation through a variety of signaling pathways (22, 27, 31, 32, 51–55). While there are many mechanisms by which lncSnhg7 promotes cell proliferation in cancer, a main feature is its ability to sponge microRNAs promoting the upregulation of downstream signaling proteins. For example, lncSnhg7 promotes pancreatic cancer proliferation through ID4 by sponging miR-342-3p (22) and promotes the proliferation of gastric cancer cells by repressing the P15 and P16 expression (53). The lncSnhg7/miR-29b/DNMT3A axis affects activation, autophagy and proliferation of hepatic stellate cells in liver fibrosis (55). Additionally, lncSnhg7 sponges miR-216b to promote proliferation and liver metastasis of colorectal cancer through upregulating GALNT1 (32) and acts as a target of miR-34a to increase GALNT7 levels and regulate the PI3K/Akt/mTOR pathway in colorectal cancer progression (31). However, the role of lncSnhg7 in normal T cell activation and proliferation is not understood.

We found that GALNT7, but not GALNT1, protein was consistently upregulated following activation of mouse CD4^+^ T cells. Overexpression of GALNT7 induces proliferation in various cancer cell lines (49, 56), but its function has not been reported in normal T cells. Recent reports have documented the importance of miR-34a in T cell activation describing it as a “hub” of T cell regulatory networks (57–59), thus it is also likely to play an important role in T cell proliferation. We found that a known target of miR-34a, GALNT7, is upregulated in stimulated T cells and that overexpression of miR-34a decreases GALNT7 protein levels. We hypothesize that the miR-34a/GALNT7 interaction is important in T cell function. However, we did observe that GALNT7 is increased in both control and Cd-exposed offspring suggesting that additional pathways may upregulate GALNT7 protein expression during T cell stimulation. Surprisingly, miR-34a overexpression did not alter GALNT7 protein levels in EL4 cells, a mouse T cell lymphoma cell line. These data indicate that T cell lymphomas may not have an intact miR-34a/GALNT7 signaling pathway. The EL4 cell line was established from a lymphoma induced in a C57BL/6 mouse by 9,10-dimethyl-1,2-benzanthracene. Chemical inductions of cancers often result in mutations which can interrupt normal signaling pathways and interactions. Alternatively, due to the inherent ability of lymphomas to replicate without external signals, the activation of the T cell receptor via CD3/CD28 may not ‘stimulate’ the cells in the same manner as primary cells.

We also found that the expression of GALNT1, another protein within the same family, is not affected by stimulation or prenatal Cd exposure. GALNT1 expression regulated by miR-216 (32). In vertebrates, expression of miR-216 is characteristic of pancreatic tissue (60) and targets of miR-216 are expressed at lower levels in pancreatic than in other tissue (61). It is not surprising that we did not observe changes in a miR-216 target as it is not highly expressed in T cells.

A limitation of our study is that neither miR-34a/GALNT7 nor miR-34a/lncSnhg7 binding was measured in the CD4^+^ T cells. We are therefore unable to directly attribute the differences in proliferation observed in the offspring of control and Cd-exposed offspring to aberrant lncSnhg7/miR-34a/GALNT7 signaling. However, we postulate that the prenatal Cd-dependent upregulation in lncSnhg7 expression induces aberrant T cell proliferation via miR-34a sequestration resulting in GALNT7 protein induction. This may occur alone or in combination with other lncSnhg7-dependent pathways.

We demonstrate that the immune alterations resulting from prenatal exposure to Cd are not due to direct exposure. Unlike other metals (e.g. lead and mercury) which transfer directly from mother to offspring, Cd concentrates in the placenta and is primarily blocked from direct transfer (1, 46, 47, 62). Our study provides the first evidence that prenatal Cd induces the expression of lncSnhg7 and alters downstream pathways in a manner that is independent of direct exposure to Cd. When we directly exposed mice to Cd, we found an induction of lncSnhg7 following T cell stimulation; however, this increase was similar in the mice exposed to control water. In contrast, the CD4^+^ T cells from mice exposed to Cd *in utero* had increased lncSnhg7 expression after stimulation as compared to mice whose parents only had access to control water. These data suggest that the immunotoxic effects prenatal Cd are induced by an indirect mechanism.

A recent study by Saintilnord *et al.* measured the effects of Cd on sperm methylation found that GALNT7 was hypermethylated in Cd exposed mice (methylation difference over control = 25.19%, p = 2.13E-18) (63). From these data we would predict that Cd-exposed males would have reduced GALNT7 expression; however, we did not observe a reduction of GALNT7 expression or reduced levels following stimulation (Figure 4). These differences may be due to cell type (somatic vs. germline) or exposure parameters (duration and dose). In our study, animals were administered 10 ppm Cd water for 21 days whereas in the Saintilnord *et al.* animals received 0.9 ppm Cd water for 9 weeks. Further studies would be needed to address these differences.

Several studies in rodents (64, 65) and humans (62) show that the toxic effects of prenatal Cd exposure may be mediated by altered zinc and copper metabolism. Zinc is essential for both innate and adaptive immune responses. It functions as an intracellular signaling molecule after T cell activation (66). However, prenatal Cd exposure is associated with zinc deficiency in the offspring (62, 64). If zinc metabolism was the mechanism driving our phenotype, we would predict decreases in lncSnhg7 and GALNT7 expression resulting in decreased proliferation. In contrast, we provide evidence that prenatal Cd exposure results in an activation phenotype in the CD4^+^ T cells. Of note, these data are consistent with stimulatory effects on T lymphocyte subsets induced by cadmium exposure previously reported by us and others (10, 67).

Cd exposure alters the activity of DNA Methyltransferases 1 and 3b (DNMT1 and DNMT3b) (68–71). In short exposures, direct Cd is an inhibitor of DNA methyltransferases and initially induces DNA hypomethylation; however, prolonged exposure results in DNA hypermethylation and enhanced DNA methyltransferase activity (71). DNA methylation is an epigenetic mechanism which may mediate the sex-specific adverse consequences of maternal Cd exposure on placental function and offspring health (72, 73). Mohanty *et al.* measured Cd-related DNA methylation in the placenta and identified sex-specific differences in the methylation of key developmental genes. Increased maternal Cd was associated with differential methylation in genes associated with cancer in placentas of female offspring. In placentas of male offspring, they found genes associated with osteoporotic fracture and kidney development were differentially methylated. Kippler *et al.* examined the associations between maternal peripheral blood and urine Cd levels and cord blood DNA methylation. Using genome-wide DNA methylation assays, they found sex-specific differences in the DNA methylation at certain genes in the offspring which correlated with higher levels of maternal Cd burden (72). They found strong associations between maternal Cd burden and differential methylation in genes related to bone morphology and mineralization and organ development in female offspring and genes related to cell death in male offspring (72). Further investigations are needed to assess the methylation patterns of lncSnhg7 within the placental and offspring of mice exposed to Cd *in utero*. This may be an indirect mechanism by which Cd exposure affects T cell function of the offspring in a sex-dependent manner.

In summary, our data demonstrate that anti-CD3/CD28 stimulation upregulates lncSnhg7 expression in CD4^+^ T cells. This effect is more apparent in the offspring exposed to Cd during gestation but is due to an indirect mechanism as direct Cd does not readily cross the placenta. CD4^+^ T cells from offspring exposed to prenatal Cd have increased proliferation when stimulated in culture. Treg suppression and apoptosis are not affected by prenatal Cd exposure suggesting that the increased proliferation is due to changes, possibly via alterations of miR-34a/GALNT7 signaling, in the CD4^+^ T cells themselves. These findings shed new light on the role of lncRNAs in T cell activation and highlight the need for understanding their functions in noncancerous and primary cells.

## 5 AUTHOR CONTRIBUTIONS

Contribution: J.L.M., C.H., and K.B. performed experiments. J.L.M., S.D., G.H., J.B.B., and I.M. analyzed results and composed figures. J.L.M, M.E.V. J.B.B., and I.M. designed experiments. J.L.M and M.E.V. composed the manuscript. All authors participated in the review of the manuscript.

## 6 CONFLICTS OF INTEREST

The authors declare no competing financial interests.

## 7 ACKNOWLEDGEMENTS

The experiments and personnel were supported by a National Institution of Environmental Health Sciences Grant ES023845 to JBB. Flow Cytometry experiments were performed in the West Virginia University Flow Cytometry & Single Cell Core Facility (RRID:SCR_017738), which is supported by the National Institutes of Health equipment grant numbers S10OD016165 and the Institutional Development Awards (IDeA) from the National Institute of General Medical Sciences of the National Institutes of Health under grant numbers P30GM121322 (TME CoBRE) and P20GM103434 (INBRE). IM was supported by West Virginia IDeA-CTR (NIH/NIGMS 2U54 GM104942-03) and National Science Foundation (NSF/1920920, NSF/1761792). The WVU Bioinformatics Core is supported by WV-INBRE grant P20 GM103434 and NIGMS Grant U54 GM-104942.

## 8 DATA AVAILABILITY

RNA-seq sequencing data were deposited in NCBI’s Gene Expression Omnibus through accession number GSE175796 (https://www.ncbi.nlm.nih.gov/geo/query/acc.cgi?acc=GSE175796).

---

**Figure S1. Ribosome-associated genes are upregulated in T cells of Cd-exposed offspring.** Gene set enrichment analysis (GSEA) of expressed genes in in vitro activated T cells from offspring against MSigDB gene set “KEGG ribosome”; genes sorted by FC of expression (parental Cd exposed/control) from high (red) to low (blue).

**Figure S2. Suppression ability of Tregs from Cd offspring is unaltered.** The suppression ability of CD4^+^CD25^+^ T regulatory cells from control and Cd offspring were assessed at 5 days.

**Table S1: List of lncRNAs differentially expressed between unstimulated and stimulated CD4+ T cells.**

## Notes

https://www.ncbi.nlm.nih.gov/geo/query/acc.cgi?acc=GSE175796

### Competing Interest Statement

The authors have declared no competing interest.

