## Supplemental Figures and Table for "Prenatal cadmium exposure alters proliferation in mouse CD4^+^ T cells via LncRNA Snhg7"

KEGG ribosome

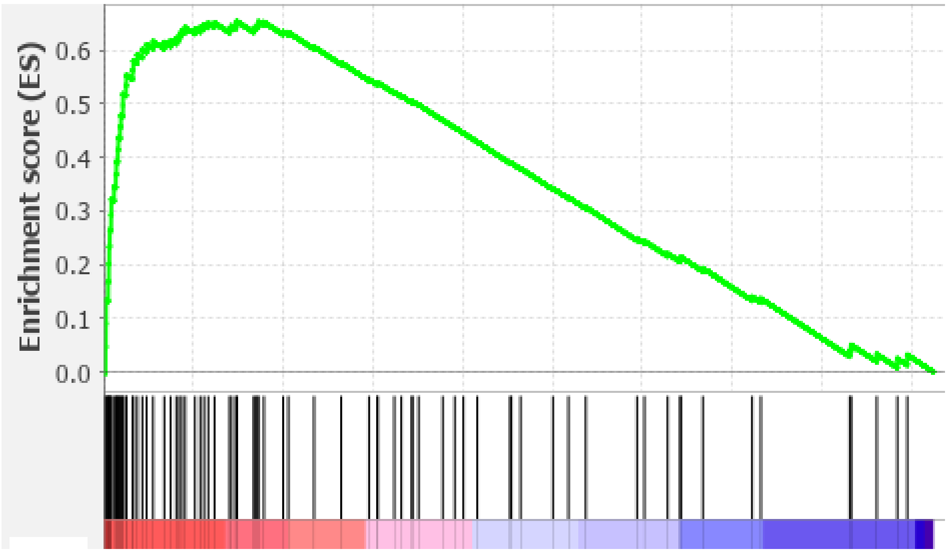


**Figure S1. Ribosome-associated genes are upregulated in T cells of Cd-exposed offspring.** Gene set enrichment analysis (GSEA) of expressed genes in in vitro activated T cells from offspring against MSigDB gene set “KEGG ribosome”; genes sorted by FC of expression (parental Cd exposed/control) from high (red) to low (blue).

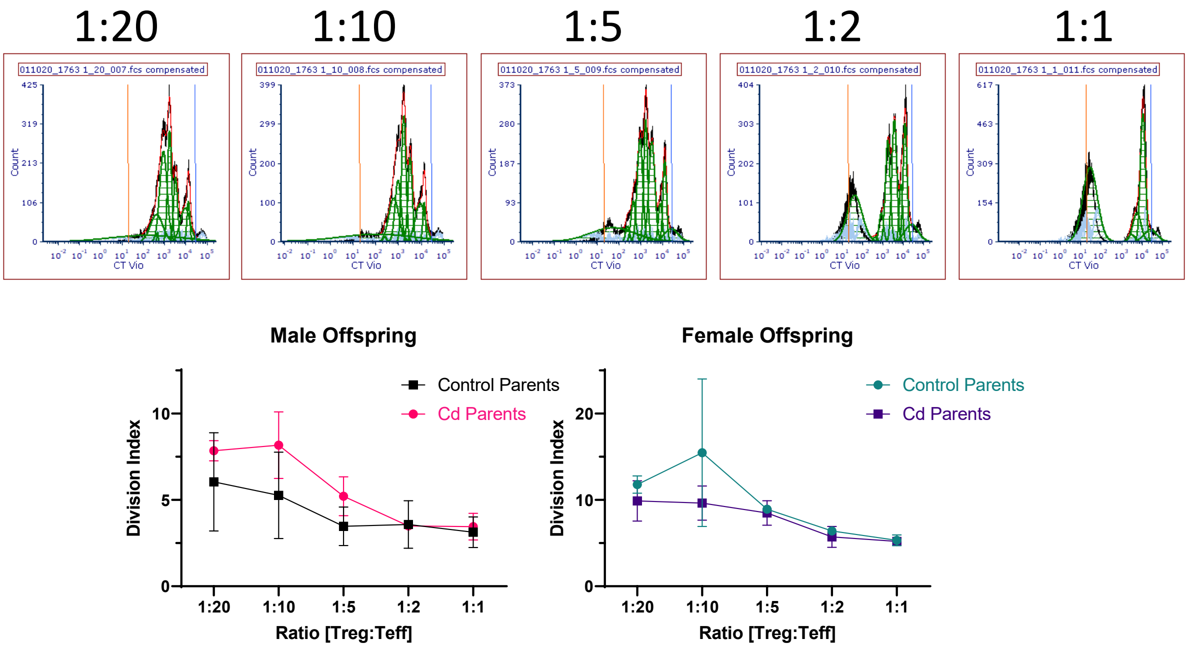


**Figure S2. Suppression ability of Tregs from Cd offspring is unaltered.** The suppression ability of $CD4^{+}CD25^{+}$ T regulatory cells from control and Cd offspring were assessed at 5 days.

| GS | logFC | FDR |
|---|---|---|
| Map2k3os | 2.98 | 1.11E-11 |
| AI506816 | 2.94 | 2.39E-44 |
| Snhg4 | 2.75 | 3.13E-72 |
| Dancr | 1.92 | 8.82E-15 |
| Snhg20 | 1.81 | 2.88E-13 |
| Rab26os | 1.71 | 5.60E-06 |
| Dnmt3aos | 1.67 | 3.20E-06 |
| Snhg7 | 1.35 | 2.27E-13 |
| Pvt1 | 1.22 | 3.05E-19 |
| AI662270 | 1.2 | 4.57E-51 |
| Smarca5-ps | 1.17 | 1.46E-29 |
| Mir17hg | 1.14 | 1.40E-26 |
| Zfp783 | 1.14 | 1.21E-12 |
| Slain1os | -1.1 | 3.58E-06 |
| H2-K2 | -1.2 | 1.24E-31 |
| Tbc1d22bos | -1.25 | 1.83E-06 |
| BC043934 | -1.3 | 1.06E-14 |
| Rapgef4os2 | -1.39 | 0.000348 |
| Zbtb11os1 | -1.39 | 2.38E-07 |
| Rbm3os | -1.41 | 4.94E-07 |
| Mir22hg | -1.63 | 7.73E-10 |
| Peg13 | -1.82 | 6.23E-105 |
| AW011738 | -1.94 | 1.56E-43 |

**Table S1: List of lncRNAs differentially expressed between unstimulated and stimulated CD4+ T cells.**
